# D- and L-lactate consumers in the human gut are taxonomically, biochemically, and energetically different

**DOI:** 10.1101/2025.10.29.685474

**Authors:** Maximilienne T. Allaart, Alexander V. Tyakht, Ruth E. Ley, Martin Pabst, Gerben R. Stouten, Largus T. Angenent

## Abstract

The human gut microbiota routinely produces D- and L-lactate during fermentation; however, the microbial fate of these stereoisomers, particularly the neurotoxic D-lactate, remains poorly understood. Given that D-lactate is an unavoidable byproduct of digestion, understanding its microbial turnover is essential for linking gut metabolism to host health. Here, we used chemostat bioreactors (pH 7.0, 37°C, and a solids retention time [SRT] of 4 d) to simulate gut-relevant, transient nutrient conditions and to enrich for lactate-consuming communities. DL-lactate-consuming consortia were enriched from a human-derived microbiota and then inoculated into duplicate bioreactors, which were fed exclusively D- or L-lactate. After steady-state was reached, the fed lactate stereoisomers were switched to assess community resilience. Regardless of the fed stereoisomer, the fermentation product spectra were consistent and dominated by acetate, propionate, and CO_2_. However, microbial communities and biomass yields diverged sharply, with a high relative abundance of *Anaerotignum* in D-lactate enrichments and *Acidipropionibacterium* and *Propionibacterium* in L-lactate enrichments. Notably, the biomass yield for D-lactate feeding was less than half that for L-lactate feeding, suggesting that the two isomers are metabolized through distinct biochemical pathways despite similar product spectra. Metagenomic and metaproteomic analyses confirmed this divergence at the pathway level. Our findings reveal how the stereoisomer identity of microbes shapes their niche specialization in the gut, with implications for understanding the ecology and clinical impact of lactate metabolism.

## Introduction

Lactate is a central intermediate in human gut fermentation and other anaerobic ecosystems [1–6]. Its conversion by specialized gut microbes produces short-chain carboxylic acids, such as propionate and *n*-butyrate, which are associated with host health benefits [7–9]. These transformations occur *via* distinct metabolic routes, including the methylmalonyl-CoA and acrylate pathways for propionate, and reverse β-oxidation for *n*-butyrate production [10]. Although these pathways are well-characterized, the ecological strategies underlying lactate consumption, especially the differential fate of its two stereoisomers, D- and L-lactate, remain poorly understood.

D-lactate, in particular, is of clinical interest. It is not produced by human cells, only within the gut microbiota, and is linked to neurodegenerative diseases, inflammation, and cancer under dysbiotic or pathophysiological conditions [11–16]. Lactate accumulation is rare in healthy individuals and balanced ecosystems, suggesting a tight coupling between lactate producers and consumers. This coupling appears across systems. In ruminant digestive tracts [12, 17], anaerobic digesters [18], soils [19], and wetlands [20], lactate remains a transient compound, only accumulating under imbalance. Despite this ecological consistency, the mechanisms by which microbial communities distinguish and metabolize lactate isomers in such environments remain unresolved.

Testing stereoisomer-specific dynamics *in vivo* is challenging due to the rapid turnover of lactate, resulting in low steady-state concentrations. Chemostat bioreactors offer a powerful alternative [21], enabling precise control of substrate input while maintaining concentrations that reflect gut-mimicking, lactate-limited conditions. These systems enable the mechanistic interrogation of microbial metabolic preferences, resilience, and ecological roles, providing an interesting platform to disentangle lactate isomer metabolism in the human gut.

The main research question we address in this study is whether D- and L-lactate are consumed by generalist or specialist microbes in the gut microbiota, and whether these stereoisomers are metabolized *via* distinct biochemical and energetic pathways. We employed chemostat bioreactors to enrich and characterize DL-lactate-consuming consortia from a humanized mouse fecal sample. Subsequently, the culture was split into bioreactors that were fed exclusively D- or L-lactate to compare fermentation stoichiometry, community composition, and growth kinetics. We identified metabolic specialization and pathway divergence between the two isomer-fed communities using metagenomic and metaproteomic analyses. Our results show that even when end products are conserved, lactate isomers select for distinct microbial taxa and strategies, underscoring the importance of stereochemistry in shaping gut microbial ecology.

## Materials and Methods

### Media composition

The cultivation of gut microbes is often performed in complex media to mimic the complexity of substrates and nutrients that are found in the human gut. The microbes residing in the gut might, therefore, rely on externally available compounds. However, using complex media makes it challenging to disentangle experimental data, because side populations can develop that feed on the complex carbon sources or proteins in gut microbe media. Here, we aimed to study lactate consumers without considerable side population growth. Therefore, we designed a rich medium based on previously developed, chemically defined media for gut microbes and for the cultivation of different types of anaerobes [22–26]. Here, we aimed to prevent the loss of lactate-consuming species with auxotrophies for vitamins or amino acids in our enrichment, while also limiting the availability of complex nutrients to prevent the development of pertinent side populations. To this end, a B vitamin solution and yeast extract were added to the medium, but they comprised only 5% of the total carbon fed to the bioreactor. The composition per liter of medium was as follows: lactate 9 g, acetate 3 g, yeast extract 0.5 g, NH_4_CL 1.5 g, KH_2_PO_4_ 0.75 g, NaCl 0.5 g, Na_2_SO_4_ 0.088 g, 1 mL trace element solution (100x), 0.1 mL vitamin solution (1000x). The trace element solution (100x) contained (g L^-1^): MgSO_4_ · 7 H_2_O 3.6; MgCl_2_ 0.2; FeSO_4_ · 7 H_2_O 0.1; CaCl_2_ 0.9; MnSO_4_ * H_2_O 0.3; ZnSO_4_ * 7 H_2_O 0.1; CoCl_2_ · 6 H_2_O 0.085; CuCl_2_ · 2 H_2_O 0.068; Na_2_MoO_4_ · 2 H_2_O 0.02; NiCl_2_ · 6H_2_O 0.02; Na_2_SeO_3_ 0.02; Na_2_WO_4_ 0.02. The vitamin solution (1000x) contained (g L^-1^): biotin (B7) 1, nicotinic acid (B3) 0.5, pABA (B10) 0.5, thiamine hydrochloride (B1) 0.5, Ca-pantothenate (B5) 0.5, pyridoxine hydrochloride (B6) 0.5, cyanocobalamin (B12) 0.1, riboflavin (B2) 0.5, folic acid (B9) 0.2, lipoic acid 0.5.

### Inoculum

A human gut microbiota-derived sample was used for inoculation of the DL-lactate enrichment. The human fecal sample [27] was used to colonize the gut of germ-free mice as described previously [28]. Mouse feces were collected after 6 weeks and stored at -80°C. Pre-cultures were prepared from a mix of male and female murine feces at 37°C in 100 mL anaerobic serum bottles containing 50 mL of the media described above but with reduced DL-lactate and acetate concentrations of 30 mM and 10 mM, respectively, and including 0.5 g L^-1^ L-cysteine and 0.5 mg L^-1^ resazurin, and N_2_ gas in the headspace. After growth was observed, the culture was passed to new serum bottles containing fresh medium. 10 mL of the grown culture from these bottles was used to inoculate Experiment I (see below).

### Bioreactor setup and control

Chemostat cultivation of lactate-consuming microbial communities, which were enriched from the human gut microbiota, was performed in 1.5-L DASGIP parallel bioreactor systems with 1.0-L working volume. The temperature was maintained at 37°C using a DASBOX4 heating block, and the pH was controlled at 7.0±0.1 using 2 M NaOH and 2 M HCl solutions, which were dosed with an MP8 pump module (DASGIP, Germany) that was controlled by custom software [29]. The bioreactors were stirred at 200 rpm using mechanical stirrers with Rushton impellers and continuously sparged with N_2_ gas at 50 mL min^-1^ using an MX4/4 module (DASGIP, Germany) to ensure anaerobic conditions. Pumping influent and effluent to and from the bioreactors was done using custom pump modules (Demo TU Delft, The Netherlands). The off-gas was cooled using a cryostat (Mini-chiller 300, Huber, Germany), which was set to 5°C to prevent evaporation of the culture broth.

### Experiment I: chemostat cultures fed with DL-lactate

To initiate the operating period of the bioreactor, the inoculum was added to 1 L of fresh medium, which was identical to that used in the anaerobic serum bottles. One bioreactor was then operated in batch mode. Considerable microbial activity was only observed after ∼30 days of incubation. Until activity was observed, the gassing conditions (headspace *vs*. liquid gassing) and stirring speed were adjusted by a trial-and-error approach to accommodate better growth. When consistent microbial activity occurred, the bioreactor was stirred at 200 rpm, and the gas was sparged through the liquid as described above. After lactate was depleted in the initial batch phase, 50% of the bioreactor volume was exchanged with fresh medium, upon which direct activity was observed. Next, lactate was depleted within 24 h after which the bioreactor was switched to chemostat mode at a dilution rate (D) of 0.01 h^-1^.

After ten days of operation in chemostat mode (*i.e.*, constant feeding and mixing), the effluent of the bioreactor was used to inoculate a second bioreactor, which was operated in identical conditions, serving as a biological replicate. Both bioreactors were initially operated at a solids retention time (SRT) of 96 h (D = 0.01 h^-1^). This retention time falls within the range of measured gut transit times in humans [30]. No considerable biofilm formation on the glass and other surfaces within the bioreactors was observed during the operation of the bioreactors. Steady state was assumed when the main product concentrations in the bioreactors and the gases in the bioreactor off-gas fluctuated <10% for at least 3 SRT periods (replicate I) or 2 SRT periods (replicate II). After maintaining steady-state conditions for 2 SRT periods, the dilution rate in the chemostat was doubled. This amounted to an SRT of 48 h (D = 0.02 h^-1^). At the end of the enrichment, the contents of the reactors were mixed and stored at 4°C. Biomass concentrations and bioreactor- and biomass-specific rates were calculated from the steady state data, and the standard deviations in these kinetic parameters were calculated using error propagation with the uncertainties package in Python.

### Experiment II: chemostat cultures fed with exclusively D- or L-lactate

Two single chemostat bioreactors that were fed exclusively with D- *or* L-lactate were inoculated with the stored broth from Experiment I, which harbored a population of DL-lactate-consuming microbes. 1 L of batch medium containing only D- or L-lactate was inoculated with 100 mL of stored broth in the two separate bioreactors. The initial batch phase took approximately ten days for D-lactate to be converted, while this took approximately four days for L-lactate. After this initial batch phase, the chemostat mode was initiated at D = 0.01 h^-1^, and the effluent from the bioreactors was used to inoculate the biological duplicates for each of the two bioreactors, resulting in four bioreactors. The operating period of the biological replicates was initiated with 500 mL of batch medium and operated in continuous mode directly. Steady state was assumed when the main product concentrations in the bioreactors and the gases in the bioreactor off-gas fluctuated <10% for at least 3 SRT periods; biological rates and their uncertainties were calculated as described for Experiment I. To calculate the statistical significance of the difference in biomass concentration between exclusive D- and L-lactate feeding, we performed a Welch’s t-test, which is suitable for systems with varying means and sample sizes.

### Analytical methods

The biomass concentration in the bioreactor was monitored with the OD_600_ by using a spectrophotometer (BioMateTM160, Thermo Scientific, United States). Cell dry weight was measured by taking 15 mL of culture broth, centrifuging, washing the pellet with MQ water, and then freeze-drying the pellet. The freeze-dried pellets were weighed, and the biomass concentration was correlated to the measured OD_600_. The resulting conversion factor from OD_600_ to biomass concentration (gDW/L) was 0.298, which was consistent among both D- and L-lactate and used for all enrichments (Supplementary Figure 5). All biomass measurements were performed in duplicate. Furthermore, the culture was inspected routinely using a microscope (Olympus BX41, Japan) to monitor for morphological changes. Samples of the culture broth were centrifuged (Eppendorf, Germany) for 5 min at 14.000 rpm, and the supernatant was filtered through a 0.22 µm PVDF syringe filter (Carl Roth, Germany). Pellets and supernatants were stored separately at -20°C until further use. Organic acids were measured from the supernatant samples with high-performance liquid chromatography (HPLC), using an Aminex HPX-87H column at 60°C coupled to a UV-VIS detector at 210 nm (Shimadzu, Germany). 5 mM sulfuric acid was used as eluent. H_2_ and CO_2_ production in the bioreactors was measured using mass spectrometry (Prima BT Benchtop, Thermo Fisher Scientific UK), with N_2_ serving as the inert gas for the gas mass balance.

### Microbial community profiling: amplicon sequencing

The microbial community composition of the bioreactors was analyzed using cell pellets from 3- or 4-mL samples taken at various time points during the enrichments. The initial microbiota composition was analyzed directly from the mouse excrement. DNA was extracted from the cell pellets or murine feces using the AllPrep PowerFecal Pro DNA/RNA extraction kit (Qiagen Inc., Germany) following the manufacturer’s instructions. The DNA content of the extracts was quantified using a Qubit 4 (Thermo Fisher Scientific, United States) to confirm that sufficient DNA was extracted for analysis. The samples were sent to Novogene Ltd. (Munich, Germany) for amplicon sequencing of the V3-V4 region of the 16S rRNA gene (position 341-806) on an Illumina NovaSeq paired-end platform; the resulting coverage was 64,000 ± 3,000 read pairs. The read processing included the following steps (software/databases): adaptor removal (cutadapt) [31]; paired merging (FLASH) [32]; quality filtering (fastp) [33]; chimera removal (vsearch) [34] using SILVA database [31, 35]); denoising and ASV generation using DADA2 [36]; and taxonomic classification of ASVs in QIIME2 [37].

For a subset of bioreactor samples, community samples from the D- and L-lactate enrichments were collected immediately before and after the substrate switch (n = 8). Shotgun metagenomic sequencing was then performed on the Illumina NovaSeq platform in 250 bp paired-end format. The genomic DNA was randomly sheared into shorter fragments, and the obtained fragments were then end-repaired, A-tailed, and further ligated with Illumina adapters. The resulting fragments with adapters were size-selected, PCR amplified, and purified. The library was quantified through Qubit and qPCR, and the size distribution was detected with a fragment analyzer. Quantified libraries were pooled and sequenced on Illumina platforms according to the effective library concentration and the required data amount. The raw reads were filtered by removing the reads containing adapters, reads with >10% undetermined bases, and reads with a low-quality score (<50% certainty of correct base calling). Shotgun metagenomic sequencing yielded 48 ± 8 Gbp paired-read coverage per sample. The sequencing data have been deposited in the European Nucleotide Archive (ENA), project accession PRJEB89830.

### Metagenomic data processing

Preprocessing of the raw metagenomic reads, assembly of metagenomes, generation of metagenome-assembled genomes (MAGs), and taxonomic classification of the MAGs were performed using an in-house pipeline [38]. Per-MAG gene prediction and functional annotation were performed using Prokka [39] and eggNOG-mapper [40]. The produced gene sequences were combined across the 102 initial MAGs to be used for the reference database in proteomic experiments. The MAGs were dereplicated at 95% ANI similarity and filtered by quality using dRep [41] to obtain 29 species-representative genomes (SRGs). These were subsequently placed into KrakenUniq [42] to prepare a reference database and quantify the species’ relative abundance in each metagenome. For the initial assessment of lactate utilization pathways that were encoded in the bacterial genomes, we prepared two lists of EC gene accessions corresponding to the essential reactions of each of the acrylate and methylmalonyl-CoA pathways for lactate utilization (n = 5 and 9 accessions, respectively). Their presence in each genome was determined based on the eggNOG-mapper-derived annotation described above. For the four MAGs representing the selected species of interest (*Anaerotignum propionicum*, *Acidipropionibacterium jensenii*, *Propionibacterium freudenreichii,* and an unclassified *Spiro-02* species), additional gene annotation was conducted using the BV-BRC portal [43] as well as the stand-alone NCBI Prokaryotic Genome Annotation Pipeline tool [44]; the three annotation tracks were subject to a fuzzy joint (allowing for a maximum of 50 bp discrepancy of the gene start and end coordinates) to form the final annotation. Generally, following the description provided previously [10], we identified lactate-utilization gene loci based on the presence of Pfam domains using hmmsearch in HMMER3. We also identified marker genes for *n*-butyrate and propionate-producing pathways and Rnf complexes *via* DIAMOND alignment [45].

### Metaproteomics

Cell pellets from 50 mL culture broth were collected from each biological duplicate after the chemostat conditions were stopped at the end of the D- and L-lactate enrichments. Protein extraction from these cell pellets and metaproteomic analysis were performed as described previously [46, 47]. Briefly, sample aliquots corresponding to approximately 250Lng of proteolytic digest were analyzed using an EASY-nLC 1200 system (Thermo Scientific, Waltham, USA) equipped with an Acclaim PepMap RSLC RP C18 column (50 µm × 150Lmm, 2 µm; Thermo Scientific, Waltham, USA) and a Q Exactive Plus Orbitrap mass spectrometer (Thermo Scientific, Germany). The flow rate was maintained at 350LnL min^-1^ with a linear gradient from 2% to 25% solvent B over 88 min, followed by an increase to 55% solvent B over 60 min. Solvent A was H_2_O with 0.1% formic acid, and solvent B was 80% acetonitrile in H_2_O with 0.1% formic acid. The Orbitrap operated in data-dependent acquisition mode, acquiring peptide signals from 385–1250Lm/z at 70K resolution, with a maximum injection time of 75Lms and an AGC target of 3*10^6^ over 175 min. The top 10 precursor ions were isolated using a 2.5Lm/z window and fragmented at a normalized collision energy (NCE) of 28. Fragment ions were acquired at 17.5K resolution, with a maximum injection time of 100Lms and an AGC target of 5e5. Singly charged ions, unassigned charges, and ions with charges >6 were excluded from fragmentation. Dynamic exclusion was set to 30 s, with peptide match preference enabled. Raw data were searched using PEAKS Studio X (Bioinformatics Solutions Inc., Canada) against protein sequences derived from assembled contigs from whole metagenome sequencing. Search parameters included a 20 ppm parent ion tolerance, 0.02Lm/z fragment ion tolerance, allowance for up to two missed cleavages, carbamidomethylation as a fixed modification, and methionine oxidation and N/Q deamidation as variable modifications. A 1% false discovery rate (FDR) was applied to peptide-spectrum matches (PSMs), and proteins were considered confidently identified if ≥2 unique peptides were matched. Label-free quantification was performed using PEAKS Q (Bioinformatics Solutions Inc., Waterloo, Canada), with peak areas normalized to the total ion current (TIC) of each run, allowing 15 ppm mass error and 7.5 min retention time variation across the dataset. Protein abundance ratios between conditions were assessed using analysis of variance (ANOVA) to determine statistical significance. Protein abundance per sample in each MAG was calculated by dividing the sample area of the protein by the total area of all proteins in that sample.

## Results

### Lactate isomer-consuming enrichment cultures grow in chemostat bioreactors

To assess the lactate stereoisomer consumption capacities of the gut microbiota, we performed two subsequent bioreactor experiments, comprising three different feeding conditions. For Experiment I, a DL-lactate-consuming microbial community was enriched from humanized mouse feces in duplicate chemostat bioreactors under gut-relevant conditions (pH 7.0, 37°C, dilution rate of 0.01 h^-1^). It took ∼30 days before considerable microbial activity was observed during the initial batch phase after which we initiated chemostat operation in the bioreactors. We characterized the steady-state fermentation parameters of the culture during Experiment I and subsequently inoculated two separate sets of duplicate bioreactors with the DL-culture, feeding them exclusively D- or L-lactate, respectively (Experiment II). Across both experiments and independently of the fed isomer, the steady-state fermentation stoichiometries were similar: acetate, propionate, and CO_2_ were consistently produced in a 1:2:1 molar ratio (Figure 1a and 3a, d, Supplementary Figure 1).

**Figure 1.**
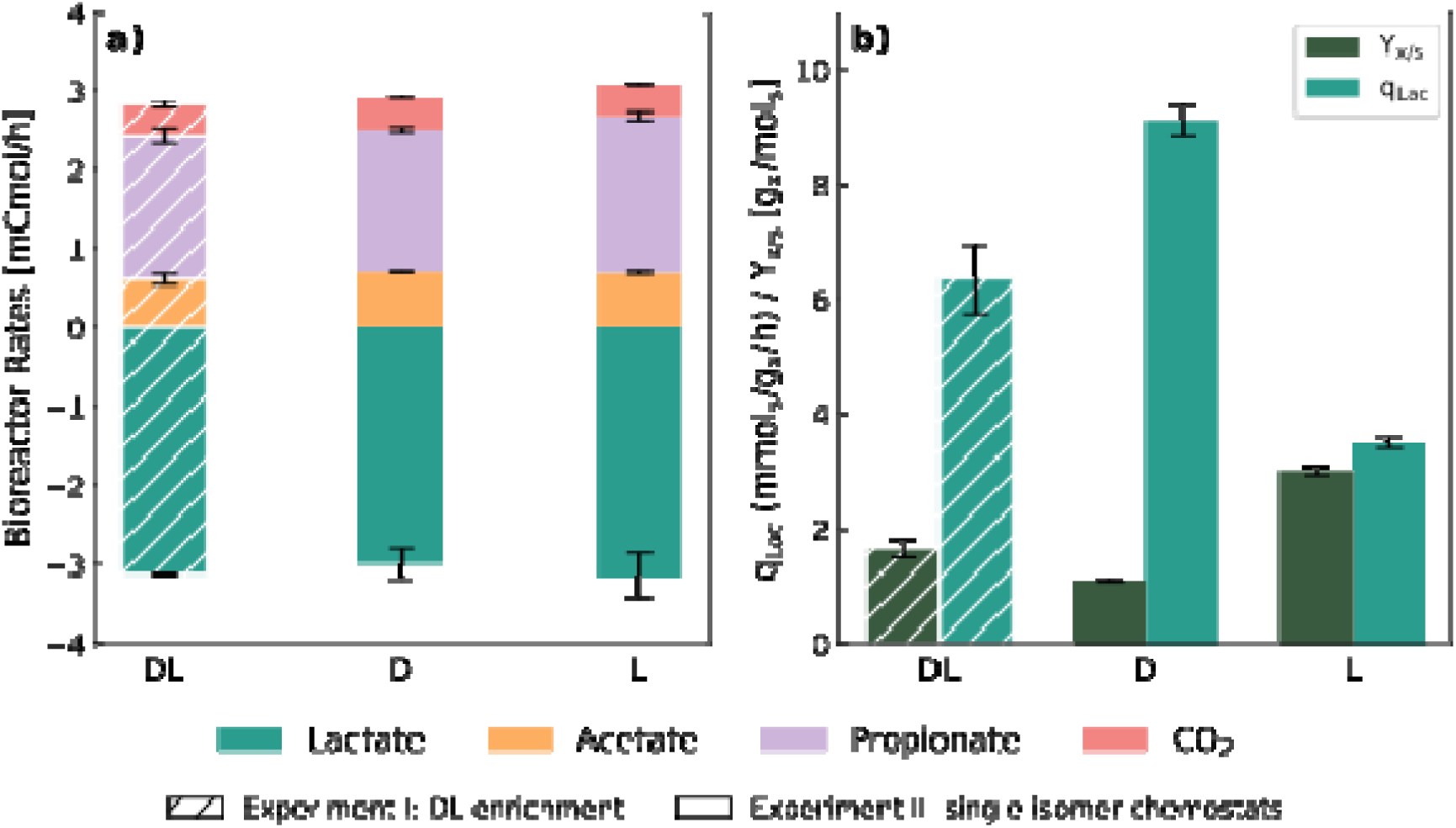
Kinetic parameters during steady-state fermentation of lactate stereoisomers in chemostats. Panel a.) shows the average (mean ± standard deviation) bioreactor-specific conversion rates of lactate and the main fermentation products during DL-(Experiment I), D-, and L-(Experiment II) lactate fermentation in steady-state chemostats in two biological duplicates (*i.e.,* n = 2 for all three conditions). This panel also represents the carbon balance, which always closed between 90 - 110% during steady-state, confirming that all main fermentation products and substrates have been identified. Panel b.) shows the average (mean ± SD) biomass yield on lactate (Y_x/s_) for each substrate, and the respective biomass-specific lactate conversion rates (q_Lac_).

Despite this metabolic convergence, community structures profiled with 16S rRNA gene amplicon sequencing diverged markedly. DL-lactate enrichments (Experiment I) were dominated by *Anaerotignum* and *Acidipropionibacterium* (Figure 2a, d). For Experiment II, D-lactate-fed chemostats selected strongly for *Anaerotignum* (Figure 2b, e), whereas L-lactate-fed chemostats initially favoured *Acidipropionibacterium* before shifting toward *Propionibacterium* dominance (Figure 2c, f). Under all conditions, *Sphaerochaeta, Bacteroides,* and *Lachnoclostridium* were part of the community, albeit with lower relative abundances compared to the main community members.

**Figure 2.**
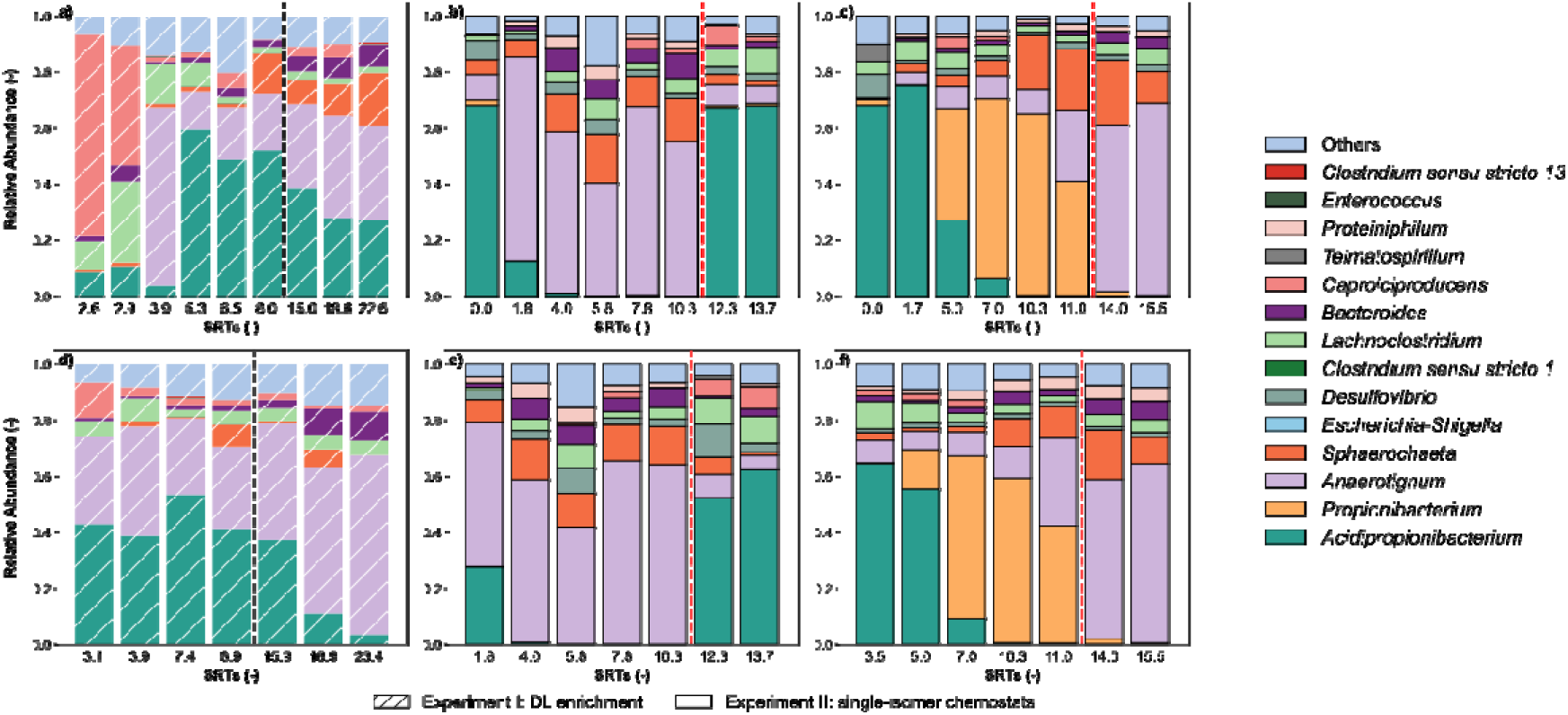
Microbial community composition development for Experiment I and II. 16S rRNA gene amplicon sequencing for community profiling in DL-lactate-fed chemostat bioreactors (Experiment I, n = 2) and exclusively D-, and L-lactate-fed bioreactors (Experiment II, n = 2 for each isomer) throughout time. Each panel represents an individual biological replicate. For Experiment I, the DL bioreactors (a, d) went through an increase in dilution rate from 0.01 h^-1^ to 0.02 h^-1^, which is indicated with the black dashed line. The SRTs at the sampling points for Experiment I were not equal because one of the bioreactors was temporarily kept at a lower dilution rate before reaching steady state to allow recovery from a technical disturbance (Supplementary Figure 1). Single-isomer fed bioreactors for Experiment II (b, e: D-lactate; c, f: L-lactate) went through a substrate switch, which is indicated with the red dashed line.

**Figure 3.**
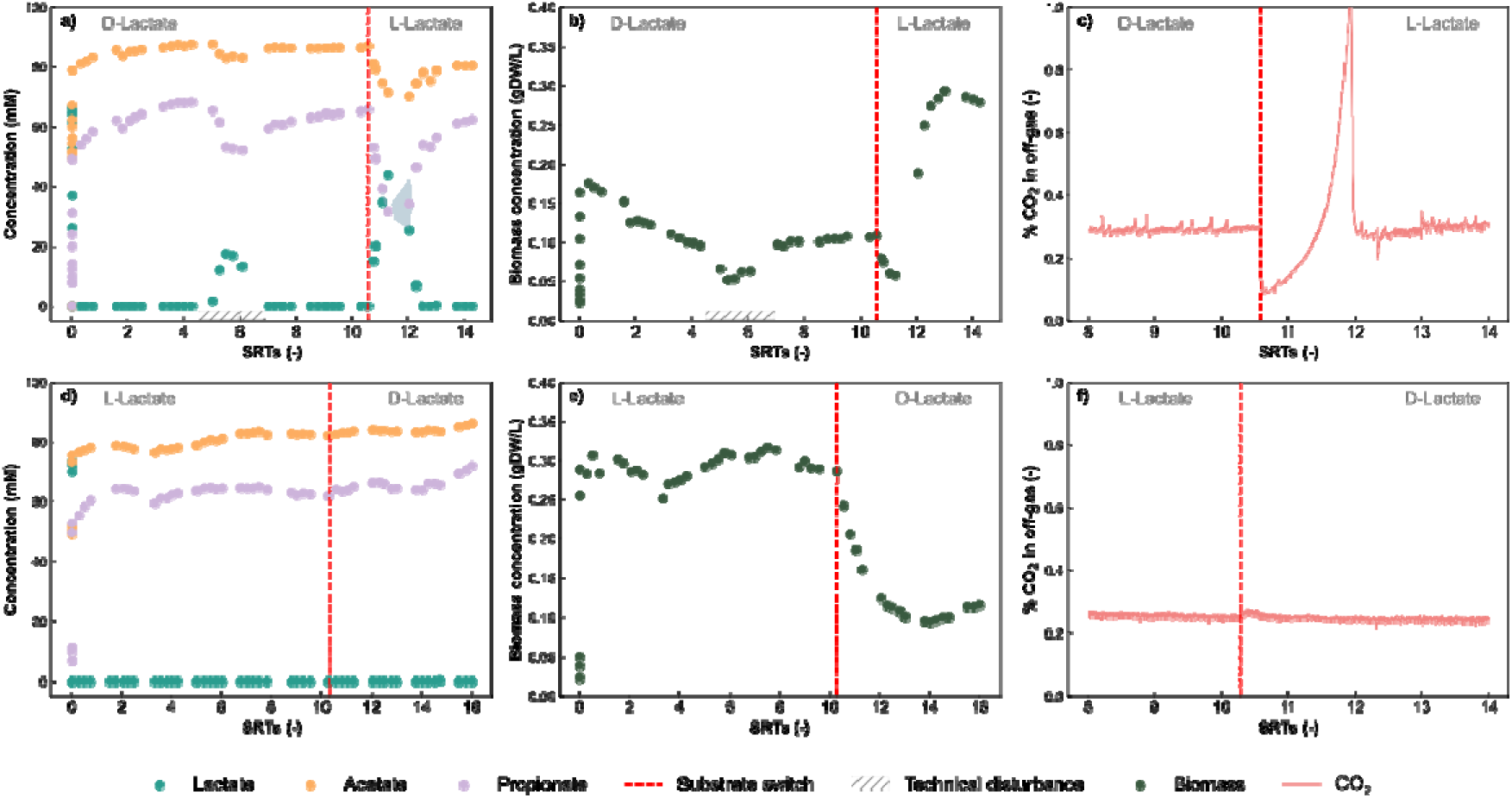
Fermentation profiles of D- and L-lactate for Experiment II chemostat bioreactors and their response to substrate switch. Panels a.) and d.) Product concentrations and panels b.) and e.) biomass concentrations in biological replicate bioreactors initially fed with D-lactate (top row, n = 2) and L-lactate (bottom row, n = 2) and switched to the other isomer, indicated with the red dashed lines. Panels c.) and f.) show the high-resolution response to the substrate switch, as observed from the CO_2_ profiles measured with off-gas mass spectrometry. Full gas profiles of all bioreactors are presented in Supplementary Figure 3. Duplicate bioreactors were started from the initial chemostat reactors after 1.5 and 2.5 SRT periods for D-lactate and L-lactate, respectively. The closed circles represent the mean of the biological replicates, and the shaded area represents the half range (*i.e.,* the distance between the measured data points and the mean). Both bioreactors initially fed with D-lactate suffered from a technical disturbance (clogged feed line) around 5 SRT periods, causing the spike in lactate concentration and dips in acetate, propionate, and biomass in panels a.) and b.), and recovered within 2 SRT periods.

Shotgun metagenomic sequencing followed by assembly and binning identified the dominant taxa as *Anaerotignum propionicum*, *Acidipropionibacterium jensenii*, *Propionibacterium freudenreichii*, *Spiro-02 sp001604275*, and *Bacteroides thetaiotaomicron* (Supplementary Figure 2). Given their origins in dairy products and the human gut, these species are likely to inhabit the human GI tract, although they are not all widely recognized as keystone gut microbial species [48–51]. This could be related to the generally low relative abundance of lactate-consuming microbes in the gut [4]. Commonly used gut microbiota composition analyses are biased towards detecting highly abundant species [52], making it entirely possible that genera or species crucial in the conversion of lactate are systematically overlooked. The enrichment approach employed here circumvents this bias.

### D- and L-lactate consumers differ in biomass yield and kinetics despite identical product spectra

To gain a deeper understanding of the physiology of the isomer-consuming microbes, we evaluated their growth kinetics. Although main product ratios were invariant, the average (mean ± SD) steady-state biomass yields differed threefold between isomers: 1.10 ± 0.03 g_x_mol_lactate_^-1^ for D-lactate *vs*. 3.01 ± 0.08 g_x_mol_lactate_^-1^ for L-lactate (Figure 1b). This discrepancy is unexpected, given identical dilution rates and input concentrations, suggesting the existence of distinct energy-conserving pathways. Biomass-specific lactate uptake rates were inversely correlated with yield: threefold higher for D-lactate consumers than for L-lactate consumers (Figure 1b). These contrasting strategies are highly significant (p<<0.001, Welch’s t-test, t = 54.6, p = 6×10^-36^; n = 18 for D-lactate, n = 20 for L-lactate) and likely alter competitive dynamics in lactate-limited systems. The affinity for substrate is a key parameter in chemostat systems, and microbes with a high affinity for the limiting substrate usually thrive. However, because the D-lactate consumers have a high biomass-specific lactate conversion rate, competition based on substrate uptake rate cannot be excluded in this system.

### Substrate switching reshapes community structure and activity

Next, we investigated the response to switching the lactate isomer in the bioreactor feed within Experiment II (Figure 3). After maintaining steady-state conditions for at least 3 SRT periods and characterizing the fermentation parameters, the lactate isomer in the bioreactor feed was switched: the D-lactate-fed bioreactors received only L-lactate, and *vice versa*. The off-gas profile (Figure 3c, f) is a real-time reflection of the microbial activity in the bioreactor [29], and the constant presence of ∼0.3% CO_2_ in the off-gas corresponded to the continuous consumption of lactate for the continuously fed bioreactors. The horizontal profiles of the figures after 2-3 SRT periods are a hallmark of steady-state chemostat operation, corresponding to stable product and biomass concentrations (Figure 3a, b, d, f).

Switching from D- to L-lactate resulted in an immediate decline in microbial activity, as evidenced by lactate accumulation and a fast decrease in the CO_2_ concentration in the off-gas (Figure 3a, c). Recovery involved exponential growth and a shift from *Anaerotignum* to *Acidipropionibacterium* dominance (Figure 3c, 2b, e). Following this exponential growth, the biomass concentration stabilized at approximately 0.28 g/L (Figure 3b), which is comparable to the biomass concentration (0.30 ± 0.008 g/L) *prior* to the substrate switch in the L-lactate enrichment (Figure 3e). Meanwhile, the concentrations of CO_2_, lactate, and product returned to steady-state values (Figure 3a). Conversely, switching from L- to D-lactate caused minor perturbation in the CO_2_ profile or product spectrum (Figure 3d, f). However, the biomass concentration gradually decreased to ± 0.13 g/L (Figure 3e), and *Propionibacterium* was replaced by *Anaerotignum* as the predominant genus.

### Combined metagenomics and proteomics reveal isomer-specific metabolic pathways

The combined physiological and community data led us to hypothesize that D- and L-lactate-consuming microbes must employ biochemically different pathways to explain the significant change in biomass concentration. To address this question, we combined differential metagenomics and metaproteomics of the community before and after the substrate switch. The relative abundance of the *Anaerotignum, Acidipropionibacterium, Propionibacterium,* and *Spiro-02* metagenome-assembled genomes (MAGs) and metaproteomes corresponded well to the relative abundance measured with 16S rRNA gene amplicon sequencing (Figure 2, Supplementary Figure 2). A single, representative high-quality MAG was identified for each of these genera. No other MAGs had a high relative abundance across all conditions.

Mapping the proteome to the metagenomic database (Figure 4) shows that the *Acidipropionibacterium jensenii* and *Propionibacterium freudenreichii* MAGs and proteomes contain the key enzymes required for the methylmalonyl-CoA pathway. *Anaerotignum propionicum*, on the other hand, does not possess the genes for the methylmalonyl-CoA pathway but instead employs the acrylate pathway for lactate utilization. *A. propionicum* was the main D-lactate-consuming taxon, and interestingly, the conversion of lactoyl-CoA to acrylyl-CoA is D-lactoyl-CoA-specific [53, 54]. This explains the isomer preference of this microbe. *Spiro-02 sp001604275* does not possess either of these pathways. Additional analysis of the lactate conversion pathways, as described previously [10], shows no involvement of *Spiro-02 sp001604275* in *n*-butyrate or 1,2-propanediol production either and confirms the use of the methylmalonyl-CoA pathway by *A. jensenii* and *P. freudenreichii* and the acrylate pathway by *A. propionicum* (Supplementary Figure 4). These two pathways are energetically distinct, with the acrylate pathway yielding one ATP per 3 mol of lactate, and the methylmalonyl-CoA pathway yielding at least 2 ^1^/_3_ ATP per 3 mol of lactate [55]. This energetic difference explains the disparity in biomass yield between D- and L-lactate (Figure 1b).

**Figure 4.**
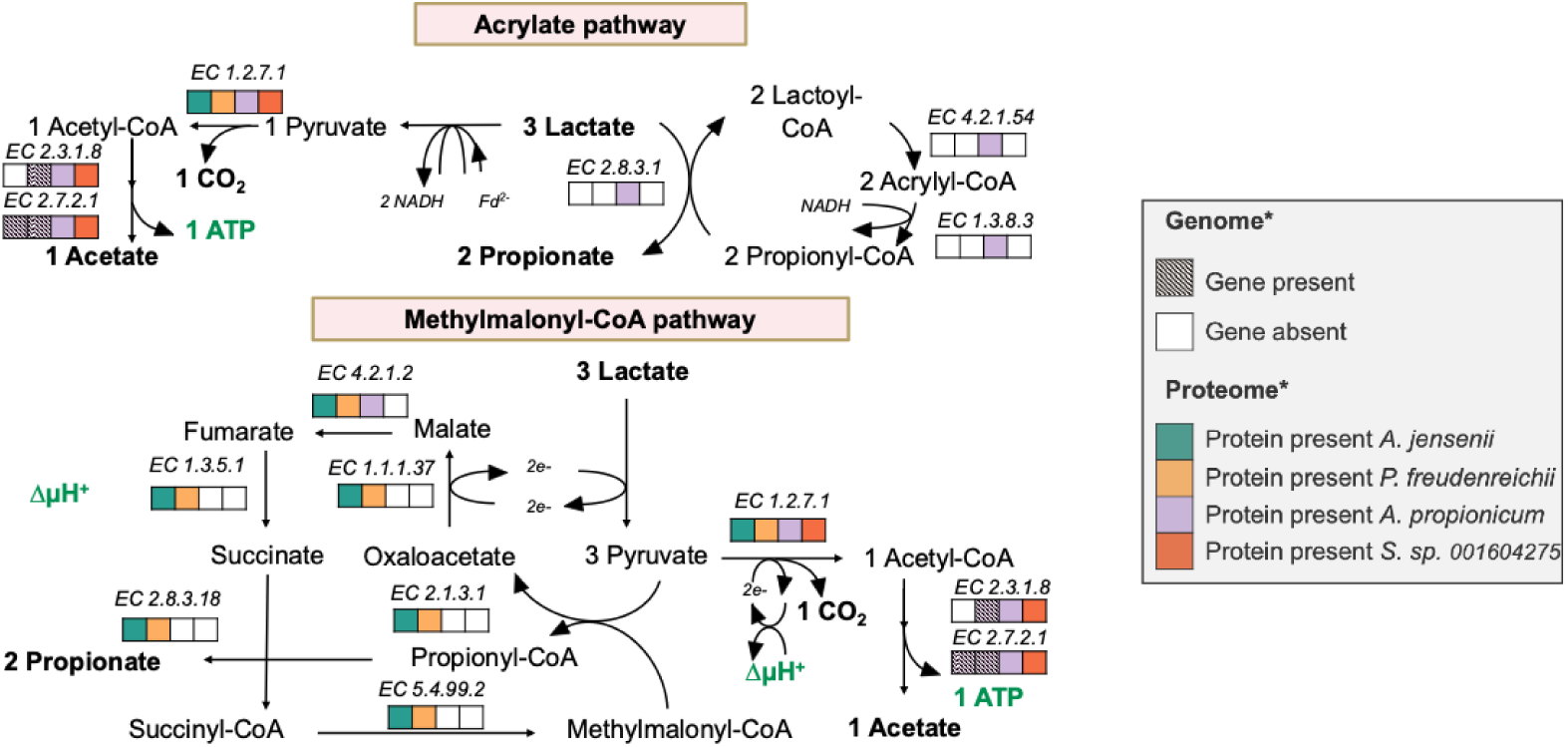
Metagenome and metaproteome analysis of Experiment II. The metaproteome of the communities was analyzed at the end of the bioreactor runs. The metagenomes provided the search database for protein identification. We identified the pathway enzymes required for the acrylate and methylmalonyl-CoA pathways based on the KEGG and Uniprot databases and separated the metaproteomics data into the proteins pertaining to our four main microbial species. Then, we searched the species’ MAGs and proteomes for the enzymes involved in the acrylate and methylmalonyl-CoA pathways. Presence of both gene and protein is indicated with a colored box, presence of the gene with a patterned grey box, and absence of both gene and protein with a white box.

### Proteomic insights into energy conservation and accessory functions

To verify the energetics of the two distinct lactate utilization pathways in our microbial communities, we utilized meta-omics data to identify the enzymes involved in lactate transport and utilization. Gene annotations of our MAGs suggest that lactate is permeated by an L-lactate permease or a glycolate transporter, which transport both D- and L-lactate despite their annotation [56]. The identification of lactate utilization genes using marker Pfam domains (see Methods) highlighted multiple loci of colocalization, providing additional evidence that many of these genes act in a co-regulated manner (Supplementary Figure 5). The cofactors used in converting lactate to pyruvate dictate the downstream pathway biochemistry, and these first enzymes are distinct in the acrylate and methylmalonyl-CoA pathways. The *lutABC* operon enables L-lactate utilization, linking it to cytochrome reduction and energy generation through the electron transport chain. Our *A. jensenii* and *P. freudenreichii* MAGs and proteomes encode these lactate utilization proteins LutABC. For D-lactate utilization, it is likely that an electron-confurcating LDH, such as LctBCD in *Acetobacterium woodie* [57], is employed by *A. propionicum.* Indeed, the EtfA and EtfB proteins (otherwise referred to as LctB and LctC) required to form the confurcating complex are abundantly present in the *A. propionicum* proteome. LctD was not found in the genome, but we identified another lactate utilization protein (FOHMDGFI_02069) [58] after refining our MAGs annotation with PGAP, which could exhibit the function of LctD. NAD-dependent D- and L-LDH (ECs 1.1.1.27 and 1.1.1.28) were also found to be encoded in the bacterial genomes from our enrichment cultures across all conditions, but have been shown to preferentially confer lactate production rather than utilization capacities [59–61].

Because the physiological profiles and meta-omics signatures for the D- and L-lactate cultures were internally consistent and clearly distinct, we examined the community proteomes for additional features that might contribute to the isomer-specific phenotypes. The most conspicuous signal was in *A. propionicum*: a single, putative tungstate ABC transporter accounted for the highest fraction of total protein; between 10-18% across samples, hinting at a tungsten-dependent redox route unique to this taxon. Furthermore, no dominant lactate consumers encoded a lactate racemase, underscoring their specialization for a single stereoisomer. Only the *Spiro-02 sp001604275* MAG carried a racemase gene, giving it the genetic capacity to interconvert D- and L-lactate; however, downstream pathway markers were either absent or below detection, so its role in lactate catabolism remains unresolved.

## Discussion

By coupling long-term chemostat cultivation with metagenomics and metaproteomics, we uncovered two functionally distinct guilds in the gut microbiota: the D-lactate specialist *A. propionicum* and L-lactate specialists *A. jensenii* and *P. freudenreichii.* These microbes are rarely identified as prevalent taxa in the human gut, suggesting that they persist at low abundance and highlighting chemostat enrichments as a powerful tool for identifying such functional groups.

Metabolic reconstruction and proteomics show that D-lactate is metabolized *via* the acrylate pathway, whereas L-lactate is metabolized *via* the methylmalonyl-CoA pathway. The import of lactate appears to be similar for both stereoisomers; however, the initial lactate-converting enzymes and their downstream biochemistry diverge, resulting in a threefold difference in biomass-specific lactate conversion rates and biomass yields between the isomers. These characteristics suggest that rate *vs*. yield trade-offs might drive the competitive strategies of lactate stereoisomer consumers [62], with D-lactate consumers gaining an advantage from high rates and L-lactate consumers gaining an advantage from high yields. The unusually large tungstate-transporting proteome fraction in *A. propionicum* supports recent reports implicating W-containing electron bi- or confurcating enzymes, hinting at potential additional energetic advantages for the D-lactate pathway [63].

The discovery of distinct biochemical pathways for D- and L-lactate conversion, as well as the absence of previously identified major gut lactate consumers (*i.e.,* microbes from the genera *Anaerobutyricum, Anaerostipes*, and *Coprococcus*), are both surprising findings. *Coprococcus* produces propionate *via* the acrylate pathway and *n*-butyrate *via* butyrate kinase [64], and *Anaerostipes* and *Anaerobutyricum* produce *n*-butyrate through reversed β-oxidation [3, 6]. The ATP yield of reversed β-oxidation is approximately 0.5 per mol lactate, which is considerably lower than the ATP yield of the methylmalonyl-CoA pathway, but not the acrylate pathway. Exploiting the methylmalonyl-CoA pathway, which we observed in the L-lactate enrichments, is the most energetically favorable scenario and is, therefore, expected in an affinity-based enrichment, such as a chemostat bioreactor. However, our data suggest that *A. propionicum* gained a competitive advantage compared to microbes with higher ATP yields by optimizing its rate, which was reflected in its high growth rate on lactate [55]. This confirms that the competition in our chemostat systems was a function of both the affinity for substrate and the substrate conversion rate. Inevitably, the operating parameters of our bioreactor systems further steered the competition, as they favored lactate specialists over multi-substrate consumers, such as the known gut lactate-consuming taxa, and might favor propionate producers due to their preference for a neutral pH [65, 66].

From the host perspective, a minority population that removes D-lactate quickly is desirable to prevent accumulation to detrimental levels. Our substrate-switch experiments demonstrated that a small population of D-lactate consumers was able to effectively cope with a rapid influx of D-lactate, which was reflected in the lack of lactate accumulation after switching from L- to D-lactate. This phenomenon was not observed for a culture enriched on D-lactate that switched to L-lactate. Whether the host actively maintains such D-lactate specialists (*e.g.*, *via* micronutrient provisioning) or whether they are sustained by cross-feeding dynamics within the microbiota remains an open question. Understanding these host-microbe interactions could inform interventions for D-lactic acidosis and potentially obesity, which was recently linked to increased circulating D-lactate levels [67].

Our data suggest that our bioreactor systems did not favor racemase-based lactate consumption. In substrate-limited chemostats, an equilibrium-based racemase [68] would halve the available concentration of the preferred isomer, and thus depress uptake rates, a probable competitive disadvantage compared to single-isomer specialists. By contrast, racemase-positive taxa are enriched in glucose-fed sequencing batch bioreactors that experience high, sustained lactate concentrations, aligning with the idea that racemization is advantageous only when substrate is abundant [69]. *Spiro-02 sp001604275* is the only racemase-encoding taxon in our communities, and the taxon does not gain dominance or encode any known lactate-converting pathways. This indicates that its growth is limited, making it unlikely that it relies solely on lactate as its primary carbon source. We, therefore, hypothesize that *Spiro-02 sp001604275* ferments protein from yeast extract, but our data is not conclusive in resolving its role in the community. A remarkable notion is the presence of *A. propionicum* with low relative abundance in the L-lactate enrichments. This observation leads to the hypothesis that *Spiro-02 sp001604275* might convert a small fraction of the L-lactate to D-lactate, thereby sustaining *A. propionicum.* In this way, a small racemase-expressing population might help sustain a population of D-lactate consumers in the human gut to maintain overcapacity for D-lactate consumption.

More broadly, nature’s strong tendency toward homochirality (*e.g.*, D-sugars in DNA and L-amino acids in proteins [70]) makes the coexistence of both lactate isomers in the gut and other fermentative ecosystems particularly intriguing. We propose that isomer-specific metabolism may confer competitive advantages to microbes that specialize in one form, and thereby enhancing ecosystem stability. On an ecosystem scale, metabolic diversity, through the presence of both D- and L-lactate-utilizing taxa, contributes to greater resilience [71] by expanding available niches. Such diversity arises not only from taxonomic variation (e.g., *Anaerotignum* vs. *Acidipropionibacterium* and *Propionibacterium*) but also from distinct metabolic pathways (acrylate *vs*. methylmalonyl-CoA) and substrate preferences (D-*vs*. L-lactate), collectively enriching the system’s functional capacity in all aspects of ecosystem diversity: species richness, genetic variation, and functional diversity [72, 73]. We further hypothesize that specialization on different isomers fosters synergistic interactions between lactate producers and consumers [74], including potential exchanges of essential amino acids or vitamins. Complementary biosynthetic capabilities and auxotrophies could reduce individual proteome burdens, leading to higher shared growth efficiencies [75, 76].

## Conclusions

Our work shows that a single human gut microbiota diverged into taxonomically, biochemically, and kinetically distinct microbial populations in D- and L-lactate-fed chemostats. These specialist taxa employed different strategies for converting their preferred lactate isomer to acetate, propionate, and CO_2_. D-lactate was converted through the acrylate pathway at higher biomass-specific rates, while L-lactate was converted through the methylmalonyl-CoA pathway, leading to a higher biomass yield. The isomer-dependent, distinct phenotype developed consistently and was replicated when the isomer in the feed was swapped. Lactate consumers comprise a small fraction of the gut microbiome, and we enriched microbial taxa that are not commonly found among lactate consumers in the gut. This suggests that cultivation-independent approaches may overlook important gut microbial species and that chemostat enrichments offer a valuable tool for studying underrepresented taxa in greater detail. The role of gut D- and L-lactate consumers in human health and disease should be further elucidated in the future

## Declarations

### Ethics approval and consent for publication

Not applicable

### Availability of data and materials

The microbiota sequencing data have been deposited in the European Nucleotide Archive (ENA), project accession PRJEB89830. The essential data analysis code is provided as a supplementary file and will be released on GitHub upon acceptance of the manuscript. Proteomics have been deposited in the PRIDE database and can be found under accession number PXD068334. (**Reviewer access details**: Reviewers can access the dataset by logging in to the PRIDE website using the following account details: **Username:**, **Password:** 4Nc64BOt8Kr8).

### Competing interests

The authors declare that they have no competing interests.

### Funding

LTA acknowledges the support from the Henriette Herz Scouting Programme of the Alexander von Humboldt Foundation to work with MTA as a post-doctoral Humboldt Research Fellow. Parts of this work were financed by grants from Cluster of Excellence - Controlling Microbes to Fight Infections (CMFI) from the University of Tübingen (EXC 2124 – 390838134).

### Author’s contributions

MTA: conceptualization, methodology, investigation, data curation, formal analysis, writing – original draft, writing – review & editing

AVT: software, formal analysis, data curation, writing – review & editing

MP: software, investigation, data curation, writing – review & editing

REL: resources, writing – review & editing

GRS: supervision, methodology, software, writing – review & editing

LTA: funding acquisition, resources, supervision, writing – review & editing

## Acknowledgments

The authors would like to gratefully acknowledge Heike Budde (Max Planck Institute of Biology, Tübingen) for supplying the inoculum of the bioreactor studies and Dita Heikens (Delft University) for metaproteomic sample preparation.

## Notes

### Competing Interest Statement

The authors have declared no competing interest.

