## Supplementary Figure for "D- and L-lactate consumers in the human gut are taxonomically, biochemically, and energetically different"

### Supplementary Information

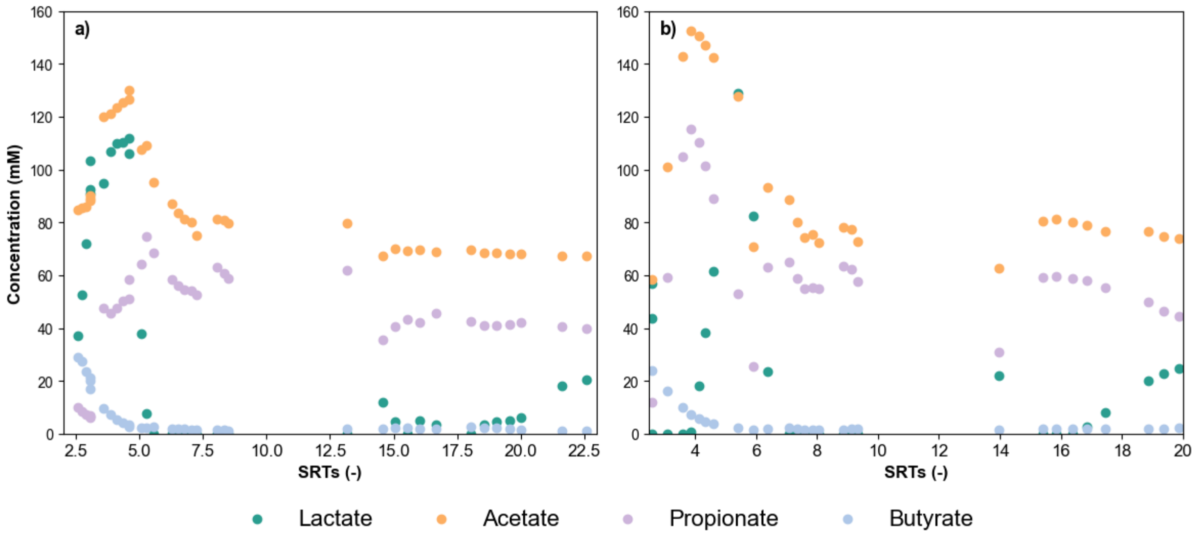


**Supplementary Figure 1. Product spectrum development in Experiment I: DL-lactate enrichment.** Panel a.) shows the product spectrum in biological replicate I. The initial start-up phase of the bioreactor was omitted from the graph. From 2.6 SRTs, the biological replicate bioreactor II, shown in panel b.), was inoculated from the effluent of replicate I. From 2.5 – 5 SRTs, the bioreactors were operated with double the amount of lactate and acetate in the feed due to a calculation error during media preparation. Initially, competition between butyrate producers and acetate/propionate producers took place, but the butyrate producers were outcompeted relatively early in the enrichment process. Due to a technical problem, a pH shock occurred after 5.5 SRTs in replicate II, after which the system required recovery. This prevented the SRTs from overlapping between replicates. The steady state at $D = 0.01\ h^{-1}$ was assumed between 5.5 and 8.5 SRTs in replicate I and 7.3 to 9.3 SRTs in replicate II, after which the dilution rate was increased to $D = 0.02\ h^{-1}$. Between 8.5 and 13 SRTs (replicate I), or 9.3 and 14 SRTs (replicate II), the bioreactor was operated without sampling. At $D = 0.02\ h^{-1}$, steady-state conditions could not be maintained because lactate concentrations increasingly accumulated.

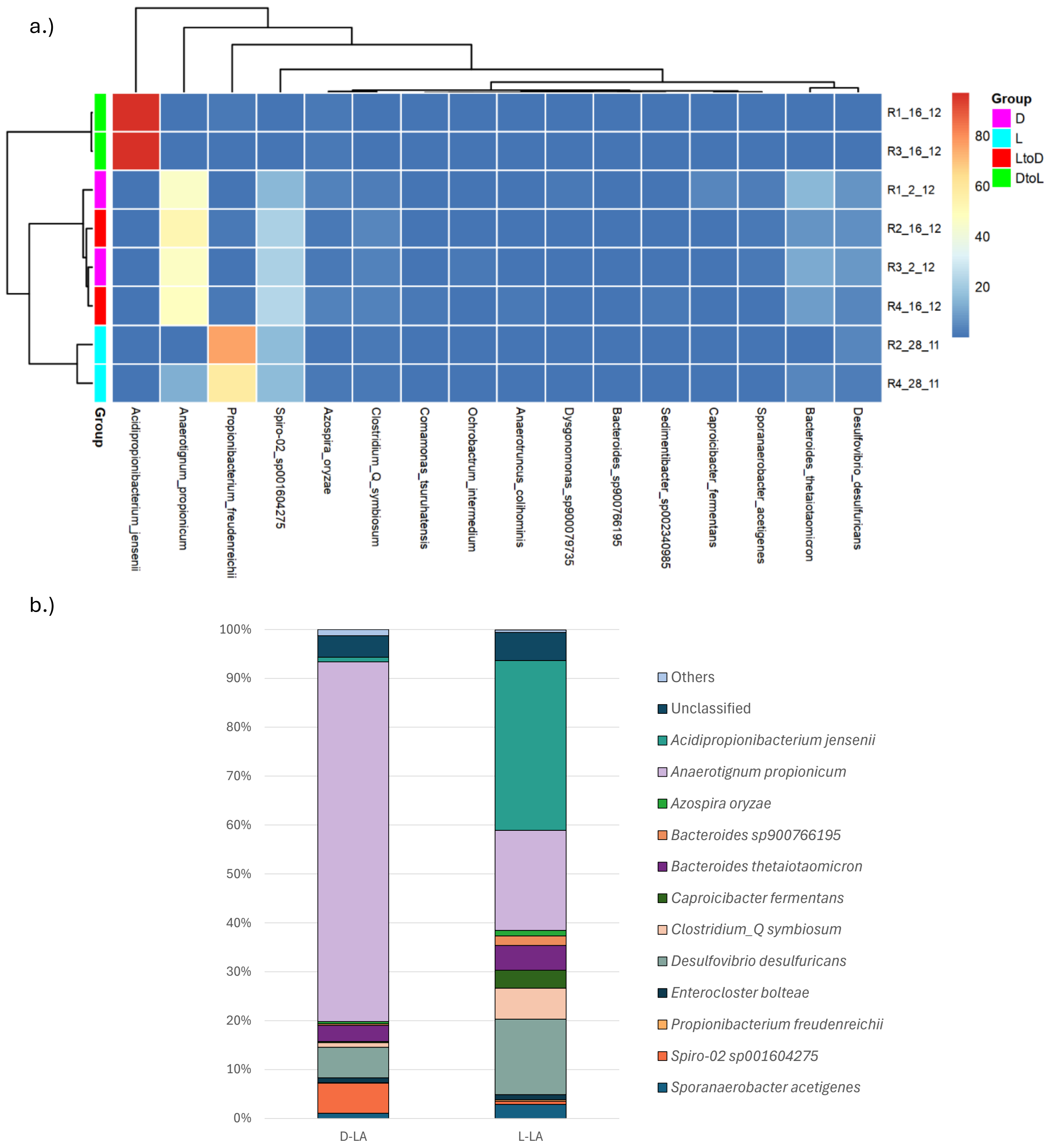


**Supplementary Figure 2. Relative abundance of the metagenomics-identified taxa in the metagenome and metaproteome.** a.) Abundance heatmap of the species identified using shotgun metagenomics. The relative abundance (%) of the species in the bioreactor samples is displayed, along with the clustering of the samples. D: sample from D-lactate-fed chemostat, L: sample from D-lactate-fed chemostat, LtoD: samples from L-lactate-fed chemostat after switching to only D-lactate feeding and a recovery period, DtoL: samples from D-lactate-fed chemostat after switching to only L-lactate feeding and a recovery period. b.) Relative abundance of each genome based on protein abundance in the metaproteome. The bar plot shows the average relative abundance of the taxa in the LtoD (left) and DtoL (right) samples. MAGs representing less than 1% of the metaproteome, except *Propionibacterium freudenreichii*, were grouped and are shown as 'Others' in the bar plot.

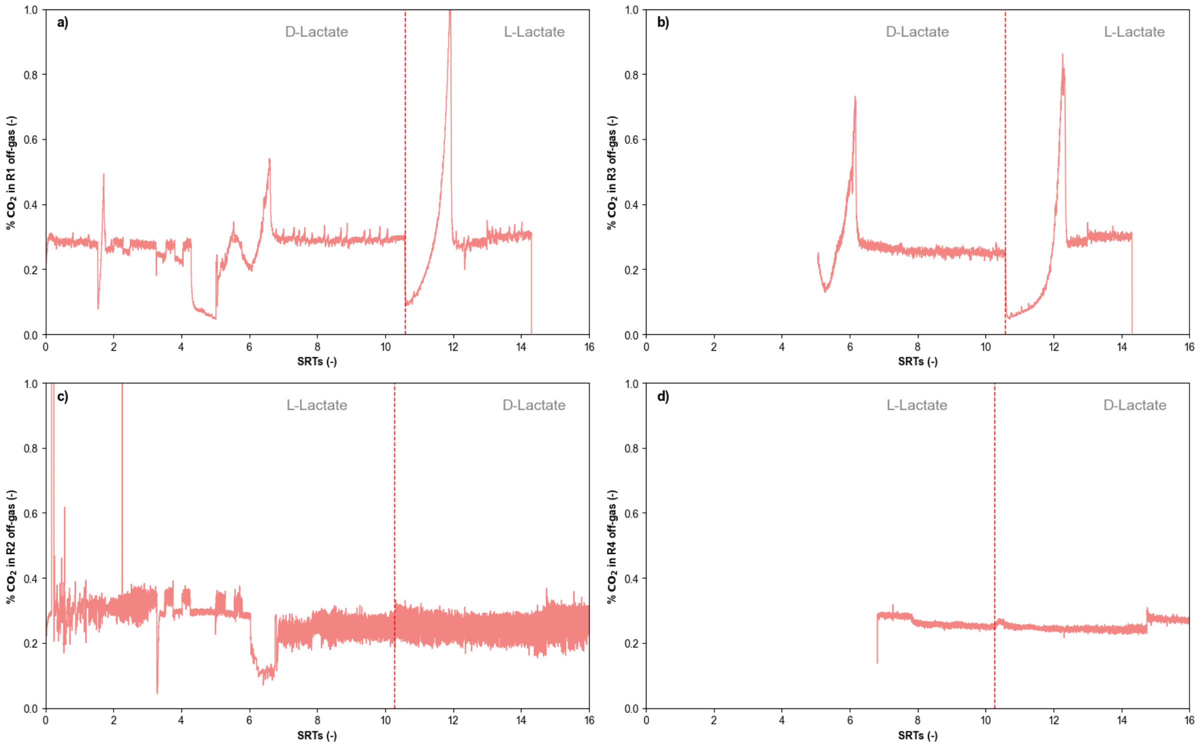


**Supplementary Figure 3. Full $CO_2$ off-gas profiles during Experiment II.** $CO_2$ profiles of all single-isomer-fed bioreactors over the entire operating period. Panels a.) and b.) show gas data of the bioreactors initially fed with D-lactate, while panels c.) and d.) show gas data of the bioreactors initially fed with L-lactate. At each measurement time, the gas stream composition was measured five times. Only the fifth measurement point was considered for these visualizations to ensure the measurement had stabilized. Off-gas measurements of bioreactors 3 and 4 (panels b. and d.) were started later during the operating period due to hardware limitations. There was consistently more noise on the channel of bioreactor 2 (panel c.), likely due to a small leak in the stream selector compartment of the mass spectrometer, causing minimal amounts of gas from other bioreactors in the lab to enter the channel. As the average concentration of $CO_2$ and $N_2$ matched the concentrations measured for the other biological replicate (panel d) and was within the same range as all other bioreactors, the data were considered reliable.

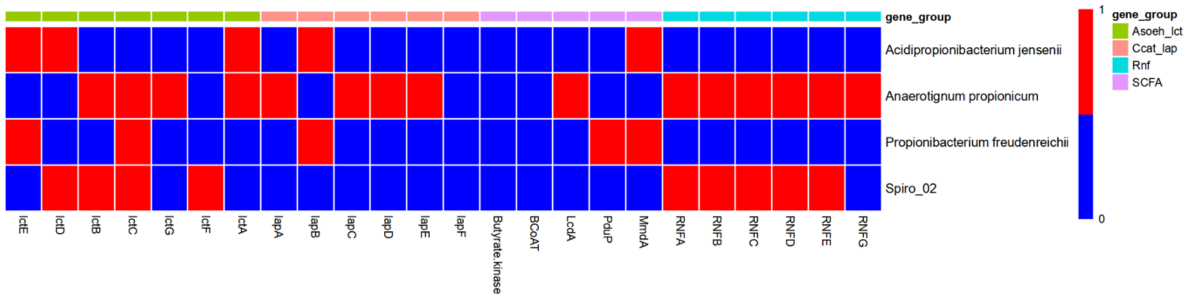


**Supplementary Figure 4. Presence of gene groups associated with lactate utilization in the selected bacterial genomes.** The genes were detected based on Diamond alignment with various thresholds of identity. Gene clusters included in the analysis were chosen according to Sheridan et al.[10] and comprised the *lct* cluster from *Anaerobutyricum soehngenii* (Asoeh_lct), the *lap* cluster from *Coprococcus catus* (Ccap_lap), Rnf complex genes (Rnf) and short-chain fatty acid production genes (SCFA). Full names of genes indicated by short-hand gene nomenclature are as follows: lactate permease (*lctE*), NAD-independent lactate dehydrogenase (*lctD*), electron transfer flavoprotein subunit beta (*lctB*), electron transfer flavoprotein subunit alpha (*lctC*), acyl-CoA dehydrogenase (*lctG*), lactate racemase (*lctF*), LutR transcriptional regulator (*lctA*), propionyl CoA transferase (*lapA*), lactoyl CoA epimerase (*lapB*), lactoyl CoA dehydratase subunits (*lapC, lapD, lapE*), lactate permease (*lapF*), butyrate:CoA transferase (*BCoAT*), lactoyl-CoA dehydratase subunit alpha (*LcdA*), CoA-dependent propionaldehyde dehydrogenase (*PduP*), methylmalonyl-CoA decarboxylase subunit alpha (*MmdAI*)*,* and Rnf complex subunits (*RNFA – G*).

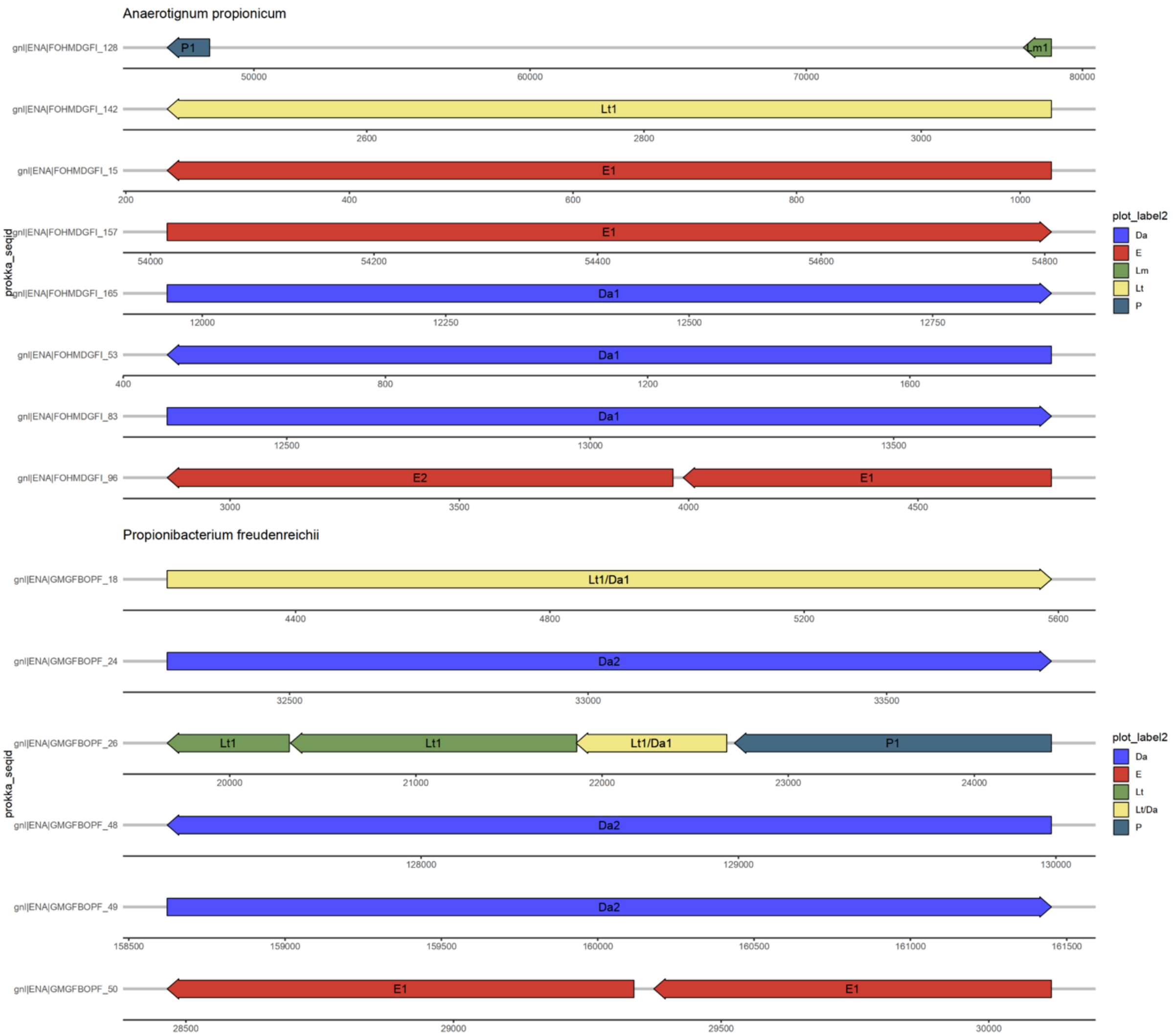
Anaerotignum propionicum
gnl|ENA|FOHMDGFI_128
P1
Lm1
50000
60000
70000
80000
gnl|ENA|FOHMDGFI_142
Lt1
2600
2800
3000
gnl|ENA|FOHMDGFI_15
E1
200
400
600
800
1000
gnl|ENA|FOHMDGFI_157
E1
54000
54200
54400
54600
54800
gnl|ENA|FOHMDGFI_165
Da1
12000
12250
12500
12750
gnl|ENA|FOHMDGFI_53
Da1
400
800
1200
1600
gnl|ENA|FOHMDGFI_83
Da1
12500
13000
13500
gnl|ENA|FOHMDGFI_96
E2
E1
3000
3500
4000
4500
prokka_seqid
plot_label2
Da
E
Lm
Lt
P
Propionibacterium freudenreichii
gnl|ENA|GMGFBOPF_18
Lt1/Da1
4400
4800
5200
5600
gnl|ENA|GMGFBOPF_24
Da2
32500
33000
33500
gnl|ENA|GMGFBOPF_26
Lt1
Lt1
Lt1/Da1
P1
20000
21000
22000
23000
24000
gnl|ENA|GMGFBOPF_48
Da2
128000
129000
130000
gnl|ENA|GMGFBOPF_49
Da2
158500
159000
159500
160000
160500
161000
161500
gnl|ENA|GMGFBOPF_50
E1
E1
28500
29000
29500
30000
prokka_seqid
plot_label2
Da
E
Lt
Lt/Da
P

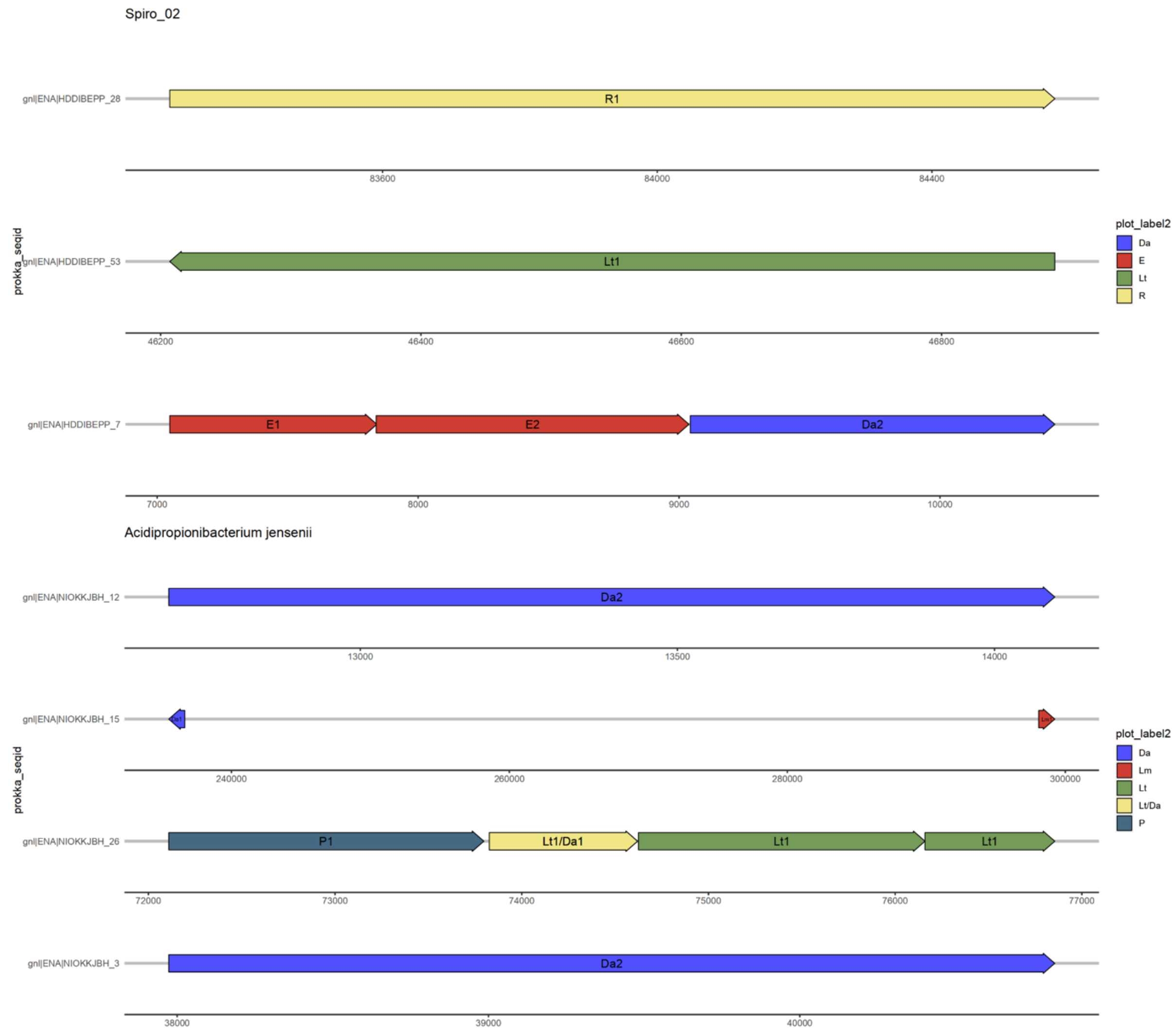


**Supplementary Figure 5. Genomic organization of lactate utilization loci.** Shown are the genes in which at least one marker functional domain of at least a lactate-utilization gene group was detected *via* hmmsearch. The labels display the name of the detected gene groups, followed by the number of respective detected domains. In cases where multiple groups are detected in a gene, these are concatenated, separated by a forward slash (“/”). Gene groups:

Lt - three-component L-lactate hydrogenase (PF02754, PF02589, PF13183, PF11870).

Da - FAD-dependent D-lactate dehydrogenase (PF01565, PF02913, PF09330, PF12838, PF02754).

Lm - FMN-dependent L-lactate dehydrogenase (PF01070, PF00173).

P - lactate permease (PF02652).

R - lactate racemase (PF09861).

E - ETF proteins alpha, beta (PF00766, PF01012).

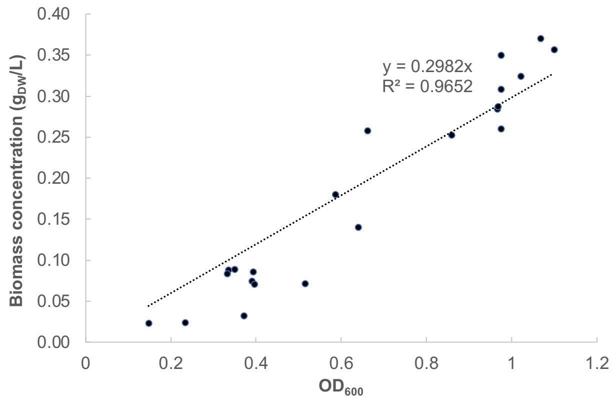


**Supplementary Figure 6. Correlation between $OD_{600}$ and broth biomass concentration.** To determine this correlation, samples from the bioreactor broth were routinely taken during the D- and L-lactate bioreactor runs. The cell dry weight was determined, and the corresponding biomass concentration was correlated to the measured $OD_{600}$ at the time of sampling. The correlation coefficient was used to convert OD measurements to biomass concentrations.
